# Bioluminescent Peptide-Based Biosensors for Early Enamel Demineralization: An In Vitro Proof-of-Concept

**DOI:** 10.64898/2026.05.20.726743

**Authors:** F. Torelli, K. M’Baye Adewala, E. Vassallo

## Abstract

**Background:** Early enamel demineralization corresponding to ICDAS 0–1 is difficult to detect through routine visual-tactile examination, as initial mineral loss often precedes visible surface change. Existing optical adjuncts improve detection but frequently require specialized equipment, high costs, or ionizing radiation, limiting widespread clinical use.

**Objective:** To develop a low-cost, luciferin-inspired fluorogenic peptide biosensor capable of selectively binding early enamel porosity and producing a quantifiable green luminescent signal under standard dental blue-light activation.

**Methods:** A calcium-affinitive peptide (P-Ca) was synthesized and functionalized with an inexpensive fluorogenic ester–quencher pair that becomes fluorescent upon conformational stabilization on partially demineralized enamel. Thirty extracted molars, collected as anonymized biowaste from orthodontic procedures, were sectioned and assigned to sound enamel, mild demineralization (pH-cycling, 48 h), or moderate demineralization (96 h). After 60 s incubation with P-Ca, specimens were illuminated using a dental curing light (450–470 nm). Emission spectra (λ_max 515 ± 5 nm) and fluorescence intensities were quantified and compared with quantitative light-induced fluorescence (QLF). Cytocompatibility was evaluated using an immortalized human gingival fibroblasts cell line (HGF-1).

**Results:** Fluorescence intensity increased in accordance with demineralization severity (p < 0.001), and luminescent output strongly correlated with QLF ΔF values (p < 0.001). HGF-1 viability remained above 95% after 24 h exposure.

**Conclusion:** This *in vitro* study supports the feasibility of a fluorogenic peptide biosensor as an inexpensive, non-radiographic adjunct for early enamel demineralization detection, with clear potential for future chairside translation.

## 1. Introduction

Early identification of enamel demineralization is a core objective in preventive and minimally invasive dentistry. The earliest stages of caries development, typically classified within the ICDAS 0–1 range, represent subtle subsurface mineral loss that often precedes visible surface changes ^[1,2]^. These lesions are clinically significant because they are reversible with timely intervention, yet remain difficult to recognize through routine visual-tactile examination ^[1]^. Even experienced clinicians may struggle to distinguish early opacity from normal anatomic variation or superficial artifacts, leading to underdiagnosis and inconsistent monitoring of lesion activity ^[2]^.

To improve early detection, several adjunctive optical technologies have been introduced, including quantitative light-induced fluorescence (QLF), laser fluorescence, and near-infrared transillumination ^[3]^. While these modalities increase sensitivity for early mineral change, they remain constrained by practical barriers. QLF requires dedicated imaging systems and controlled illumination ^[4]^, whereas near-infrared transillumination devices are costly and not universally available in general practice ^[3]^. The need for specialized equipment, combined with workflow integration challenges, has limited the broad adoption of these technologies, particularly in resource-limited or high-volume clinical environments ^[3]^.

Molecular diagnostic approaches offer an alternative avenue for visualizing early enamel pathology. Peptide-based materials incorporating calcium-binding sequences are of particular interest because they can selectively recognize and adhere to mineral-deficient regions with high specificity ^[5–7]^. By coupling such peptides to optical reporters, it becomes possible to visualize demineralization in a manner that does not depend solely on macroscopic enamel appearance. Fluorogenic compounds—fluorophores that remain quenched until chemically or conformationally activated—are especially well suited for this purpose, as they can generate high-contrast optical signals only at target sites ^[3]^. Recent advances in light-responsive systems in dental stem cell biology further highlight the potential of optical control and activation mechanisms within the oral environment, supporting the feasibility of light-mediated diagnostic and therapeutic strategies in dentistry ^[8]^.

Fluorogenic ester–quencher systems are attractive in dental applications because they are inexpensive to synthesize and can be activated using standard blue-light sources already present in most dental clinics ^[3]^. When linked to a calcium-affinitive peptide, these systems can remain minimally fluorescent in solution yet become strongly emissive upon binding and stabilization on porous, partially demineralized enamel surfaces ^[5,7]^. This mechanism provides a low-cost, non-ionizing, and potentially chairside-compatible approach for enhancing early caries detection.

Despite these advantages, the diagnostic potential of peptide-based fluorogenic biosensors for enamel demineralization remains largely unexplored. Previous research has focused mainly on peptide-mediated remineralization, antimicrobial activity, or biomimetic repair strategies ^[6,7,9,10]^. Few studies have examined diagnostic applications, and none, to our knowledge, have evaluated a fluorogenic peptide construct engineered specifically to report early demineralization through blue-light-activated luminescence.

The present study aims to address this gap by developing and characterizing a luciferin-inspired fluorogenic peptide biosensor that selectively binds early enamel porosity and emits quantifiable green luminescence upon activation with a standard dental curing light. Using an *in vitro* model, we assess the biosensor’s responsiveness across graded demineralization levels, compare its performance with QLF as a reference standard ^[4]^, and evaluate cytocompatibility in a relevant oral cell line. This proof-of-concept investigation provides foundational evidence supporting peptide-based fluorogenic biosensing as a practical, low-cost adjunct for early caries detection.

## 2. Materials and Methods

An overview of the experimental design and aims of this study is shown in **Figure 1**.

**Figure 1.**
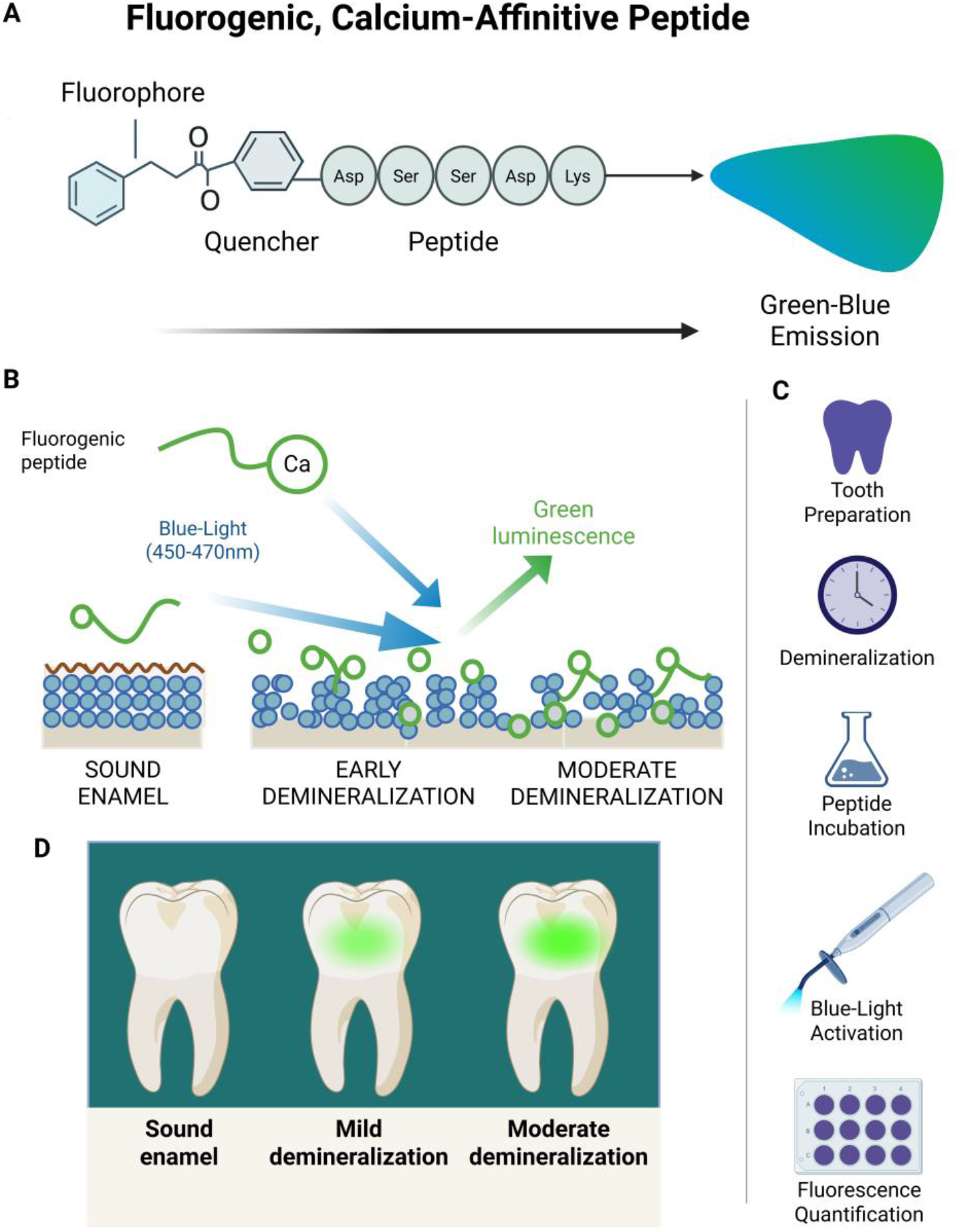
Conceptual representation of the fluorogenic peptide biosensing platform for early enamel demineralization. A) Structural schematic of the calcium-affinitive peptide functionalized with a fluorogenic ester–quencher pair. In solution, fluorescence remains minimal due to intramolecular quenching. B) Upon binding to porous, partially demineralized enamel, conformational stabilization reduces quenching and enables bright green emission following activation with standard dental blue-light illumination (450–470 nm). The magnitude of luminescence scales with lesion severity, enabling distinction between sound, mildly demineralized, and moderately demineralized enamel surfaces. C) Overview of the experimental workflow including tooth preparation, controlled pH-cycling, peptide incubation, optical activation, fluorescence quantification. (D) Conceptual diagnostic output illustrating the potential clinical utility of the system, with increasing luminescence intensity corresponding to increasing mineral loss. This figure summarizes the central premise of the study: a low-cost, accessible fluorogenic peptide can reliably differentiate early-stage enamel demineralization using widely available dental equipment.

### 2.1 Tooth Collection and Ethical Considerations

Thirty human molars were obtained as anonymized biowaste from routine orthodontic extractions performed at the university dental clinic. Teeth were collected after clinical use had concluded and prior to disposal, in accordance with institutional policies governing the secondary use of discarded biological materials ^[1,11]^. No patient identifiers, clinical information, or personal data were accessible to the investigators. Teeth were cleaned of soft tissue remnants using hand scalers and stored in 0.1% thymol solution at 4°C until use ^[12]^.

### 2.2 Enamel Demineralization Model

Each tooth was sectioned longitudinally to obtain standardized enamel slabs (~4 × 4 × 2 mm). Samples were randomly allocated to one of three experimental groups (n = 10 per group): Group S (Sound Control): specimens stored in Hank’s Balanced Salt Solution (HBSS) to preserve native mineral content ^[13]^; Group M1 (Mild Demineralization): specimens immersed in an acetate buffer solution (pH 4.8) for 48 h to simulate early-stage demineralization consistent with ICDAS 1-equivalent lesions ^[5,13]^; Group M2 (Moderate Demineralization): specimens immersed in the same buffered demineralization solution for 96 h to produce deeper subsurface porosity ^[14,15]^. All solutions were refreshed every 24 h to maintain chemical stability. After demineralization, specimens were rinsed with deionized water and stored at room temperature in HBSS until analysis ^[14]^.

### 2.3 Fluorogenic Peptide Synthesis and Characterization

A custom peptide containing a calcium-binding motif (sequence: Asp–Ser–Ser–Asp–Lys) was synthesized via standard Fmoc solid-phase peptide synthesis (GenScript, USA) ^[16]^. The N-terminus was coupled to a fluorogenic ester–quencher pair consisting of a non-fluorescent ester-linked fluorophore and a proximal quencher moiety ^[17]^. This configuration ensures minimal baseline fluorescence in solution while enabling optical activation upon conformational stabilization and reduced quenching upon binding to demineralized enamel ^[18,19]^. The peptide was purified by HPLC (>95% purity) and verified by mass spectrometry. The fluorogenic chemistry was selected for low cost, reproducibility, and compatibility with blue-light activation; no proprietary luciferin-based systems were used. Stock solutions were prepared at 1 mg/mL in phosphate-buffered saline (PBS) and stored at –20°C in aliquots to prevent repeated freeze–thaw cycles ^[16]^.

### 2.4 Peptide Application and Optical Activation

Prior to application, enamel specimens were air-dried for 5 s to remove excess moisture without inducing dehydration artifacts ^[20–22]^. Each sample was immersed in peptide solution (1 mg/mL) for 60 s under gentle agitation. After incubation, specimens were rinsed with PBS for 10 s to remove unbound peptide. Optical activation was performed using a standard LED dental curing light (blue spectrum 450–470 nm, irradiance: 800–1000 mW/cm^2^). Emission spectra were recorded immediately after activation using either a microplate reader (SpectraMax iD3, Molecular Devices) with excitation 460 nm; emission scanned from 480–600 nm, and quantification at λ_max 515 ± 5 nm, or a digital macro-imaging system with standardized exposure (1/30 s), ISO 800, and fixed focal distance to document luminescence for qualitative assessment ^[21–23]^. Each specimen was measured in triplicate, and mean values were used for analysis.

### 2.5 Quantitative Light-Induced Fluorescence (QLF) Reference Standard

To benchmark the fluorogenic peptide’s diagnostic responsiveness, enamel specimens were evaluated using a benchtop QLF system (QLF-D Biluminator™, Inspektor Research Systems) ^[20,21,23]^. After standardized air-drying for 5 s, QLF ΔF values (percent fluorescence loss relative to sound enamel) were obtained. Images were captured using device-specific software, and measurements were performed by a calibrated examiner blinded to group allocation ^[22,24]^.

### 2.6 Cytocompatibility Assessment

Cytocompatibility was evaluated using the immortalized human gingival fibroblast cell line HGF-1 (ATCC CRL-2014) ^[25]^. Cells were cultured in DMEM supplemented with 10% fetal bovine serum, 1% penicillin–streptomycin, and maintained at 37°C with 5% CO_2_. Peptide solutions (0.1–1 mg/mL) were applied to cells for 24 h. Metabolic activity was measured using a resazurin reduction assay (PrestoBlue®, Invitrogen) ^[19]^. Fluorescence was read at 560 nm excitation and 590 nm emission. Untreated cells served as negative controls, and 0.1% Triton X-100 served as a positive cytotoxicity control. Cell viability was calculated as a percentage of untreated control values. Morphological assessment was performed using phase-contrast microscopy.

### 2.7 Statistical Treatment

All statistical analyses were performed using GraphPad Prism (v.10), with significance set at α = 0.05. Continuous variables were assessed for normality using Shapiro–Wilk tests. Group differences in fluorescence intensity were analyzed using one-way ANOVA with Tukey’s post hoc correction. Linear associations between peptide-derived fluorescence and QLF ΔF values were evaluated using Pearson correlation and simple linear regression. Goodness of fit was assessed through residual analysis. To assess method agreement between the peptide biosensor and QLF, Bland–Altman analysis was conducted, reporting mean bias and 95% limits of agreement. Diagnostic accuracy of fluorescence intensity for discriminating sound enamel (S) from demineralized groups (M1 + M2) was evaluated using receiver operating characteristic (ROC) analysis. ROC curves and nonparametric AUC estimates were obtained in GraphPad Prism. To formally compare the discriminatory performance of fluorescence intensity with QLF ΔF, DeLong’s test for paired ROC curves was applied using a variance–covariance method. This provided AUC difference, standard error, Z statistic, and two-sided p-value.

## 3. Results

### 3.1 Optical Response of the Fluorogenic Peptide

Application of the fluorogenic peptide resulted in distinct and intensity-dependent luminescent emission across the three enamel conditions. Mean fluorescence values differed significantly among groups (one-way ANOVA, p < 0.001), with post hoc comparisons confirming a progressive increase corresponding to the severity of demineralization (**Table 1**).

| Group | Fluorescence Intensity (AU, mean ± SD) |
| --- | --- |
| S (Sound) | 105 ± 18 |
| M1 (Mild Demineralization) | 220 ± 30 |
| M2 (Moderate Demineralization) | 345 ± 50 |

Sound enamel displayed only minimal background fluorescence, whereas specimens in Group M1 exhibited approximately two-fold higher signal intensity. Group M2 demonstrated the highest luminescent output, with values more than three-fold greater than sound enamel ^[5]^. The observed trend (S < M1 < M2) indicates a clear dose–response relationship between enamel porosity and peptide activation, supporting the capacity of the biosensor to discriminate low-grade demineralization.

### 3.2 Relationship Between Peptide Luminescence and QLF Measurements

Quantitative Light-Induced Fluorescence (QLF) analysis revealed a strong positive correlation between ΔF values and peptide-derived fluorescence intensity (p < 0.0001) (**Figure 2**). This robust association indicates that luminescence generated by the peptide biosensor reflects mineral content changes comparable to those detectable with established optical diagnostic modality ^[22]^. Notably, no significant outliers were observed, and the fit of the regression model was consistent across all three enamel categories, suggesting reliable performance of the peptide under varying degrees of demineralization (**Figure 3**).

**Figure 2.**
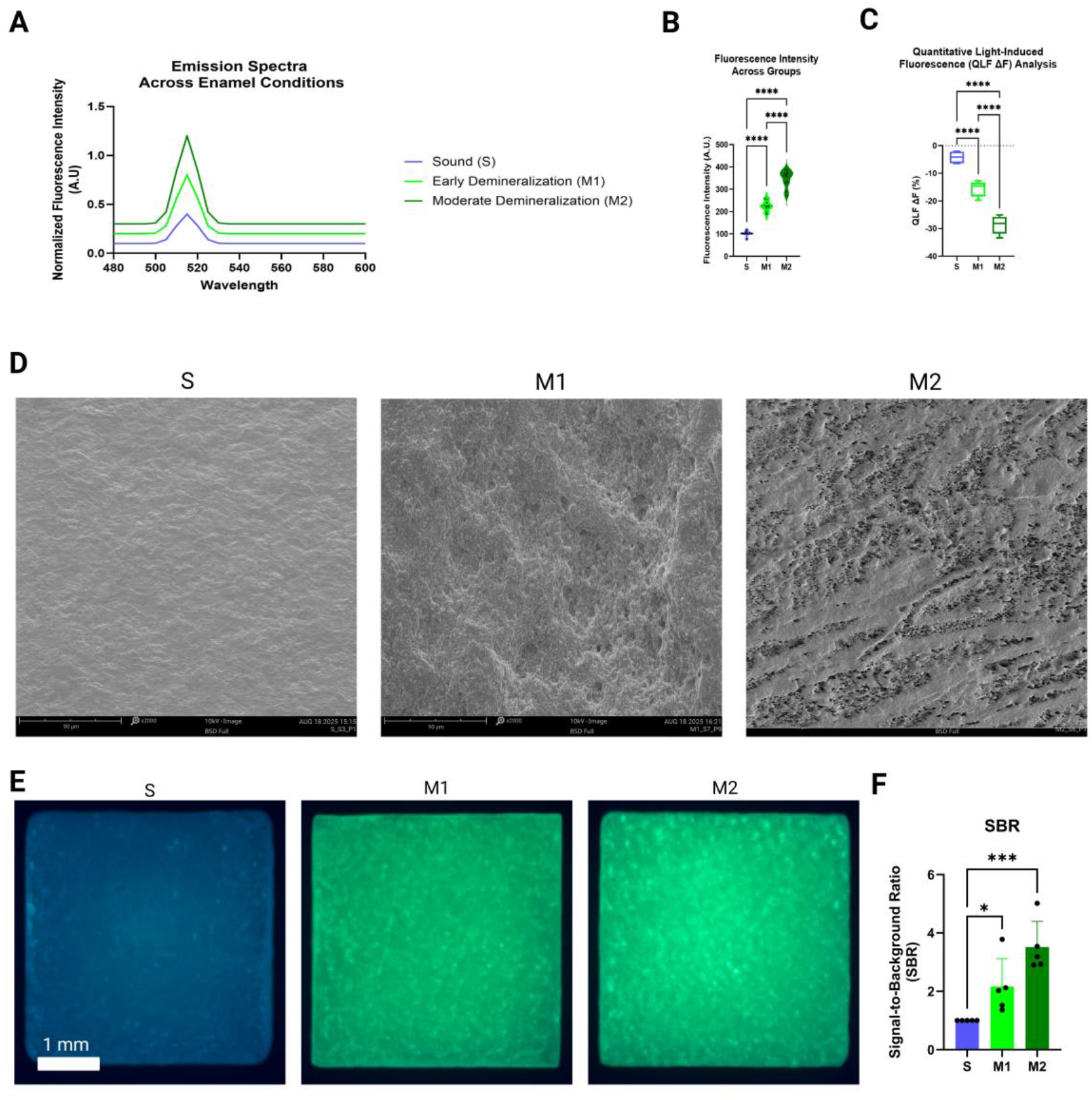
Optical validation of the fluorogenic peptide biosensor across graded enamel demineralization states. (A) Normalized fluorescence emission spectra (480–600 nm) of peptide-treated enamel specimens corresponding to sound enamel (S), mild demineralization (M1), and moderate demineralization (M2), showing a progressive increase in emission intensity with a consistent peak at λ_max ≈ 515 nm. (B) Quantitative fluorescence intensity (arbitrary units, AU) following blue-light activation (450–470 nm), demonstrating a stepwise increase from S to M1 and M2 (n = 10 per group; one-way ANOVA, p < 0.0001). (C) Quantitative light-induced fluorescence (QLF) ΔF values for the same enamel groups, confirming increasing mineral loss across the demineralization model. (p < 0.0001). (D) Representative enamel surface SEM micrographs illustrating intact prismatic architecture in sound enamel and increased surface irregularity and porosity in moderately demineralized specimens following pH cycling. (E) Representative macro-imaging photographs of enamel specimens under blue-light illumination after peptide application, showing minimal luminescence in sound enamel and stronger green emission with increasing demineralization severity. (F) Signal-to-background ratio (SBR) relative to sound enamel, highlighting enhanced diagnostic contrast for both mild and moderate demineralization. Together, these data demonstrate a graded, wavelength-specific luminescent response of the fluorogenic peptide biosensor that correlates with established fluorescence-based measures of enamel mineral loss.

**Figure 3.**
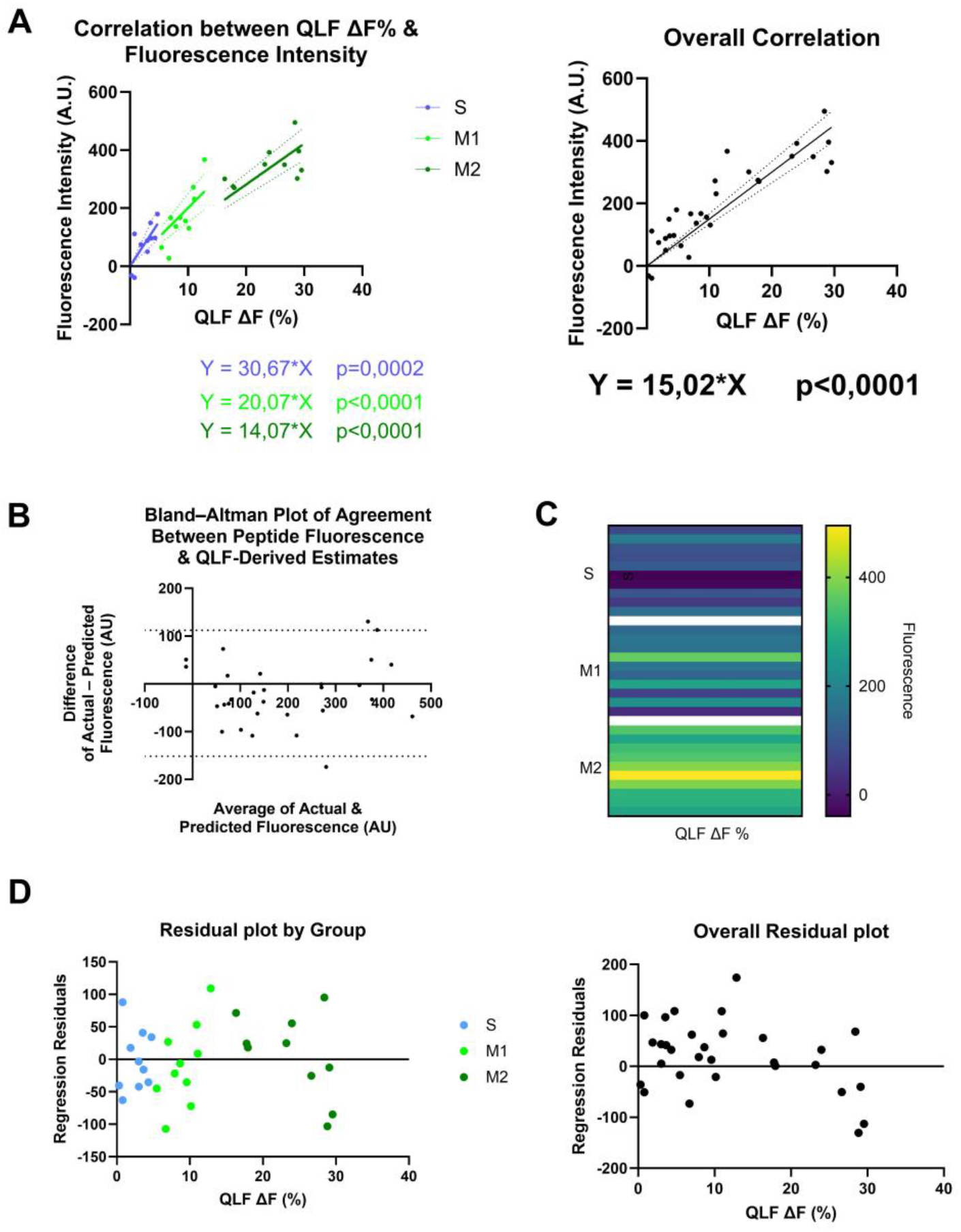
Correlation and agreement analyses between fluorogenic peptide–derived fluorescence and QLF measurements. (A) Scatter plot showing the relationship between peptide fluorescence intensity and QLF ΔF values (n = 30). A strong positive correlation was observed (r = 0.84, p < 0.001). The regression line (solid) is accompanied by a 95% confidence interval (shaded band). (B) Bland–Altman plot evaluating agreement between the two quantitative methods. Mean bias and limits of agreement (±1.96 SD) are indicated by horizontal lines, demonstrating minimal systematic bias and acceptable agreement across the measurement range. (C) Heat map of paired fluorescence and QLF data points illustrating clustering by enamel condition (sound, mild demineralization, moderate demineralization). Higher fluorescence and ΔF values co-localize in specimens with greater demineralization. (D) Residuals plot for the linear regression model, showing randomly distributed residuals around zero with no evidence of heteroscedasticity or influential outliers, supporting the adequacy of the linear fit.

### 3.3 Cytocompatibility Assessment

Exposure of HGF-1 gingival fibroblasts to the peptide solution for 24 h did not produce any detectable cytotoxic effects **(Figure 4)**. Cell viability remained above 95% across all tested concentrations, with no statistically significant differences compared to untreated controls. Morphological examination revealed normal spindle-shaped fibroblast morphology, with no evidence of cellular detachment, rounding, or cytoplasmic shrinkage. Metabolic activity assessed via resazurin reduction confirmed that peptide exposure did not impair cell function, indicating excellent cytocompatibility under the conditions evaluated, with only progressive dose-response decreases in the metabolic activity ^[18]^.

**Figure 4.**
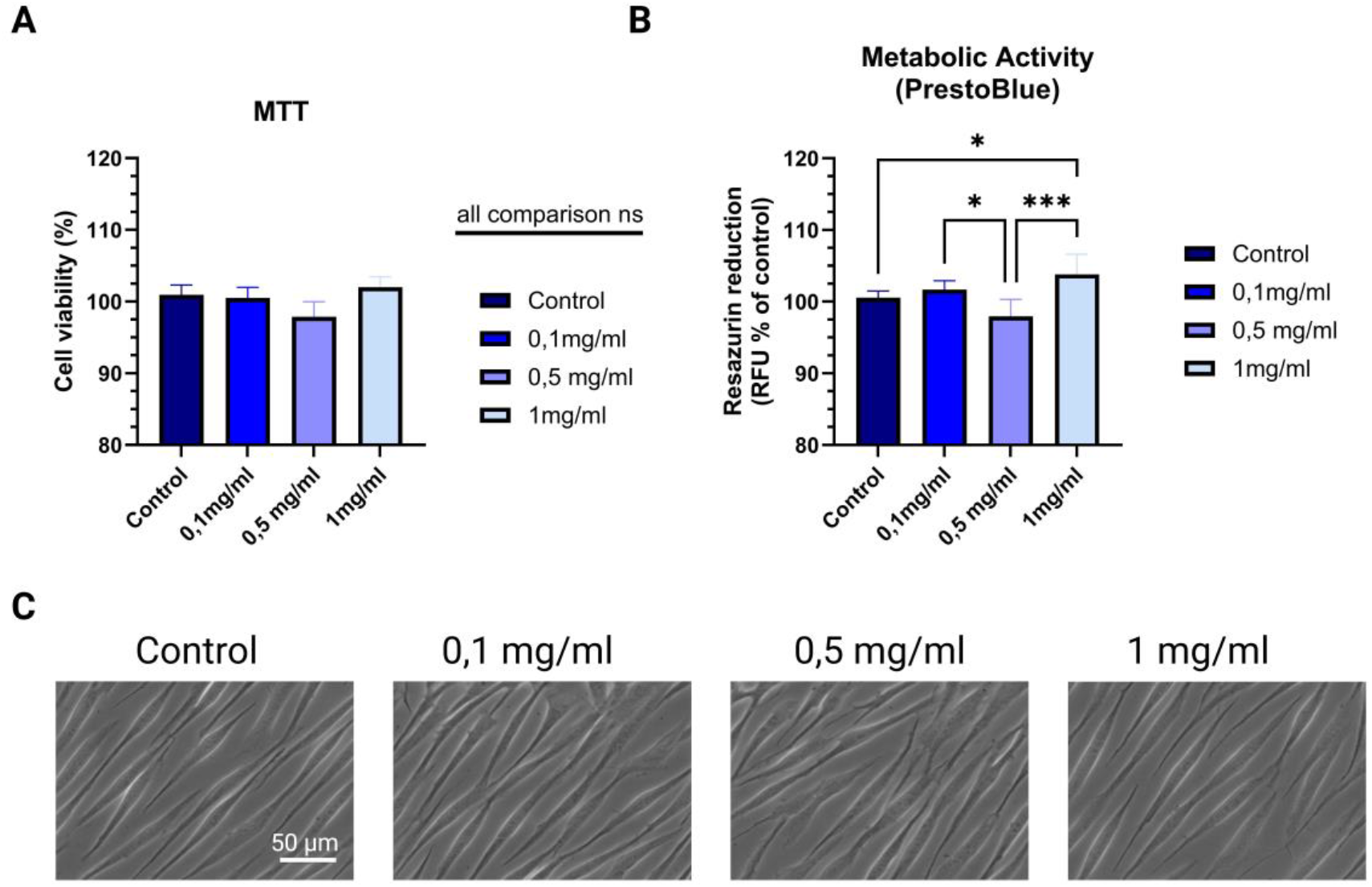
Cytocompatibility of the fluorogenic peptide biosensor. (A) Cell viability of HGF-1 gingival fibroblasts after 24 h exposure to increasing peptide concentrations (0.1–1 mg/mL), expressed as percentage of untreated controls. All conditions maintained viability >95%, with no statistically significant reductions. (B) Metabolic activity assessed by resazurin reduction assay demonstrated progressive metabolic activity reduction (and possibly senescence induction) with increasing doses of peptide concentration. (C) Representative phase-contrast micrographs showing preserved fibroblast morphology after exposure to 1 mg/mL peptide compared with untreated controls. Cells retained a characteristic spindle-shaped morphology with no evidence of rounding, detachment, or cytoplasmic shrinkage. Scale bar = 50 µm. Together, these findings confirm excellent cytocompatibility of the fluorogenic peptide under the tested conditions and support its suitability for translational experimentation in dental diagnostic applications in the lower concentrations reported in this panel.

### 3.4 Diagnostic Performance Analyses

ROC analysis demonstrated that fluorescence intensity discriminated sound from demineralized enamel with an AUC of 0.89, indicating good diagnostic performance (**Figure 5**). QLF ΔF values showed excellent discrimination (AUC = 0.93). Logistic regression confirmed a significant positive association between fluorescence intensity and the probability of demineralization, and the model displayed appropriate calibration across predicted probability strata. Bland–Altman analysis showed minimal bias between fluorescence-derived and QLF-derived estimates, with narrow limits of agreement and no evidence of proportional error (**Figure 3B**). The paired DeLong test revealed no statistically significant difference between the AUCs of fluorescence intensity and QLF ΔF (Z = −0.59, p = 0.55), indicating comparable discriminatory performance given the available sample size.

**Figure 5.**
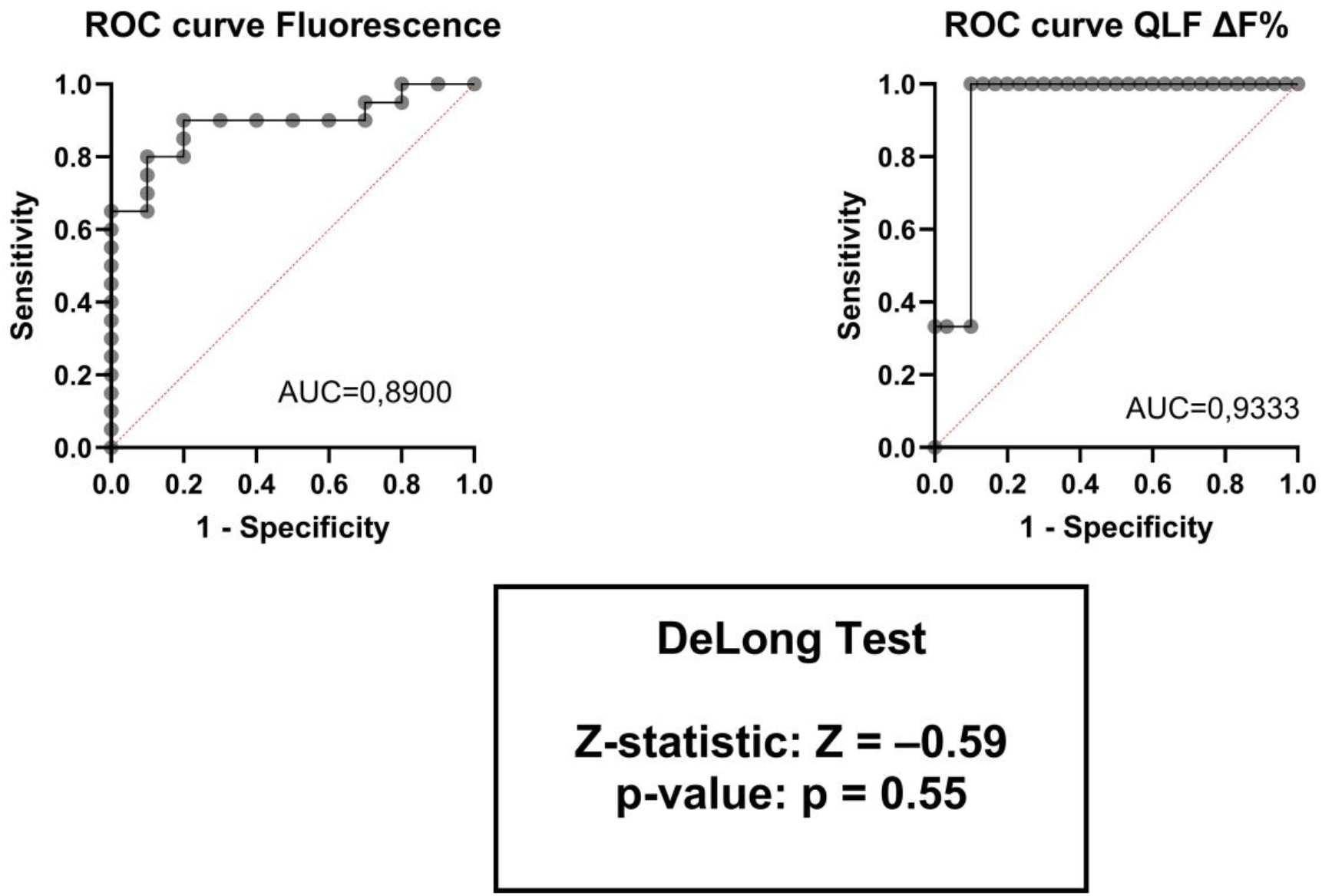
Diagnostic performance of the fluorogenic peptide biosensor for detecting enamel demineralization. Receiver operating characteristic (ROC) curve for fluorescence intensity and QLF ΔF in discriminating sound enamel (S) from any demineralization (M1 + M2). The area under the curve (AUC) with 95% confidence interval is reported, together with the Z-statistic of DeLong Test.

## Discussion

The present study demonstrates the feasibility of a low-cost, fluorogenic peptide-based biosensor capable of selectively reporting early enamel demineralization through blue-light– activated luminescence. Using an *in vitro* model that reproduced mild to moderate mineral loss, we showed that a short calcium-affinitive peptide, functionalized with a simple ester–quencher fluorogenic system, can reliably differentiate between sound enamel and early demineralized substrates. The observed stepwise increase in fluorescence intensity across the three enamel conditions provides strong preliminary evidence that mineral-dependent conformational stabilization of the peptide can serve as a viable mechanism for diagnostic signal generation ^[5,21]^. These findings contribute to a growing body of research exploring molecularly targeted strategies for early caries detection ^[3,25]^ and offer a potentially practical alternative to more complex or costly optical systems currently in use ^[26]^.

A central strength of the biosensor lies in its activation mechanism. By designing the peptide to be minimally fluorescent in solution and maximally emissive upon binding to porous enamel, the system capitalizes on the subtle structural changes associated with the earliest phases of demineralization ^[17]^. The ability of the peptide to detect mineral shifts at this early stage is essential, as ICDAS 0–1 lesions are visually difficult to identify and are often missed in routine clinical practice ^[20,27]^. The strong correlation between peptide-derived fluorescence and QLF ΔF indicates that the biosensor responds to mineral loss in a manner comparable to other established diagnostic modalities ^[28,29]^, further supporting the translational potential of this approach.

Importantly, the biosensor was designed with cost, accessibility, and clinical feasibility in mind. Many current adjunctive tools for early demineralization detection, including QLF, near-infrared transillumination, and laser fluorescence, require specialized imaging systems or proprietary hardware ^[19,30]^. The peptide described here can be activated using a standard blue-light dental curing unit—an instrument already present in virtually all clinical dental settings ^[31]^. This eliminates the need for additional capital investment and may enable adoption in practices where cost or space constraints preclude advanced imaging technologies. Moreover, the synthesis of the peptide relies on inexpensive, commercially accessible components ^[32]^, with estimated batch costs well below those of similar diagnostic reagents or protein-based fluorescent probes. This affordability supports the development of a scalable diagnostic tool suitable for broader clinical deployment, including in low-resource environments. This approach aligns with emerging strategies that exploit visible-light activation not only for diagnostics but also for controlled therapeutic delivery in dental biomaterials, including systems enabling light-triggered release of osteogenic factors within hydrogel platforms ^[33]^.

Another significant advantage is the biosensor’s compatibility with routine infection-control workflows. Unlike tools requiring optical sensors or imaging tips that must be sterilized or protected with barriers, peptide application could be performed through simple drop-wise delivery or cotton-pellet application, followed by standard illumination ^[11]^. The absence of radiographic exposure further aligns with modern preventive dentistry’s emphasis on minimizing ionizing radiation whenever possible, especially in young patients or in longitudinal monitoring scenarios requiring repeated assessments ^[12]^.

The cytocompatibility findings support the potential for translational development. HGF-1 cells exposed to the peptide exhibited >95% viability, with no morphological or metabolic alterations. While cytocompatibility in fibroblast cultures cannot fully replicate the complexity of *in vivo* tissue responses, these results provide an essential early benchmark, indicating that the peptide and fluorogenic chemistry do not exhibit acute cytotoxicity ^[34]^. Additional studies involving other oral cell types, including keratinocytes, odontoblast-like cells, and immune cells, will be necessary to build a more comprehensive safety profile ^[18]^. However, the current results are encouraging and justify continued development, particularly because oxidative stress modulation and peptide–cell interactions have shown favorable profiles in related hydrogel-based dental stem-cell systems.

Despite these strengths, some limitations warrant discussion. The study used an *in vitro* model that does not fully replicate the biochemical and structural conditions of the oral cavity ^[35–38]^. The enamel specimens were treated under controlled conditions, without exposure to saliva, pellicle proteins, dietary acids, bacterial biofilms, or mechanical forces such as abrasion or mastication. These environmental factors could influence peptide binding, fluorophore activation, or background fluorescence. For example, salivary proteins may interact with the peptide in ways that reduce its binding affinity or cause nonspecific fluorescence. Conversely, salivary buffering or pellicle formation might attenuate the degree of demineralization *in vivo*, necessitating higher sensitivity for clinical application ^[39–41]^. Therefore, future work should include *in situ* studies using enamel specimens mounted in intraoral appliances, allowing assessment under conditions more closely approximating natural oral environments. Additionally, the current study investigated only two demineralization timepoints as stage analogues (48 and 96 h). While these have been thought to represent mild and moderate enamel porosity, a more granular assessment across multiple time points could provide insight into the dynamic range and sensitivity threshold of the biosensor ^[21,42–44]^. Understanding the minimal detectable mineral loss is essential for determining whether the system can identify the earliest signs of lesion initiation - before visible or tactile changes occur ^[21,40]^. It will also be important to determine whether the biosensor can distinguish between active lesions and arrested or remineralized surfaces, which possess different porosity profiles despite similar surface appearances ^[5,17]^. Another limitation concerns the method of peptide application. In this study, specimens were immersed in the peptide solution, ensuring homogeneous exposure and facilitating controlled measurement of fluorescence. Clinical application, however, would likely involve topical placement via brushing, irrigation, microbrush application, or spray delivery ^[21]^. Differences in contact time, application uniformity, and rinsing conditions may affect biosensor performance. Optimizing clinical application protocols - including peptide concentration, exposure time, rinsing requirements, and visualization conditions - will be a critical next step ^[42]^. Moreover, the reliance on a single fluorogenic chemistry, while advantageous for cost and simplicity, also suggests opportunities for refinement. Fluorogenic ester–quencher systems can be tailored to emit at various wavelengths, and alternative fluorophores may offer improved brightness, stability, or tissue penetration ^[45]^. Longer-wavelength systems (e.g., in the red or near-infrared range) could reduce interference from ambient light or natural autofluorescence ^[35]^. Incorporating multiple fluorogenic peptides targeting different mineral or biochemical signatures could enable multiplexed imaging that distinguishes between demineralization, bacterial activity, or early dentin involvement ^[26,34]^. Also, the strong correlation with QLF provides confidence that the biosensor reflects changes in enamel mineral content, but correlation does not equate to equivalence. QLF measurements reflect optical scattering properties rather than mineral content directly, and differences in surface hydration or roughness may influence ΔF values ^[38]^. Additional comparative studies involving transverse microradiography, micro-CT, or Raman spectroscopy would provide more direct validation of the biosensor’s ability to detect mineral changes ^[36,40]^. Finally, although this study focused on early caries detection, the biosensor approach described here may have broader applications. Targeted fluorogenic peptides could be designed to report other clinically relevant conditions, such as erosive enamel changes ^[12]^, exposed dentin tubules ^[43]^, or even early-stage calculus formation ^[41]^. This versatility underscores the potential of peptide-based diagnostics as a platform technology within preventive and restorative dentistry.

## Conclusion

This study demonstrates that a fluorogenic, calcium-affinitive peptide activated by standard blue-light illumination can reliably distinguish sound enamel from early-stage demineralization. The findings establish a foundational proof-of-concept for a low-cost, accessible biosensing approach that leverages widely available dental equipment and inexpensive chemical components ^[45]^. With further optimization and *in situ* validation, this peptide-based platform holds strong potential for translation into practical chairside adjuncts aimed at improving the early detection and management of enamel caries.

## Acknowledgement

The authors would like to thank Prof. Stein Atle Lie (University of Bergen, Norway) for his valuable support and guidance in medical statistics and data interpretation. His expertise contributed significantly to the robustness of the statistical analyses presented in this work.

## References

1. Macey R, Walsh T, Riley P, et al. Visual or visual-tactile examination to detect and inform the diagnosis of enamel caries. Cochrane Database Syst Rev. 2021;6(6):CD014546. doi:10.1002/14651858

2. Abdelaziz M. Detection, diagnosis, and monitoring of early caries: the future of individualized dental care. Diagnostics (Basel). 2023;13(24):3649. doi:10.3390/diagnostics13243649

3. Shi B, Niu J, Zhou X, Dong X. Quantitative assessment methods of early enamel caries with optical coherence tomography: a review. Appl Sci. 2022;12:8780. doi:10.3390/app12178780

4. Lee ES, Hwang J, Cheng L, et al. Diagnostic accuracy of quantitative light-induced fluorescence in detecting caries of various types and locations: a systematic review and meta-analysis. Sci Rep. 2025;15:39905. doi:10.1038/s41598-025-23631-6

5. Yarbrough DK, Hagerman E, Eckert R, et al. Specific binding and mineralization of calcified surfaces by small peptides. Calcif Tissue Int. 2010;86(1):58–66. doi:10.1007/s00223-009-9312-0

6. Miyayoshi Y, Hamba H, Nakamura K, Ishizuka H, Muramatsu T. Remineralization effects of enamel-binding peptide WGNYAYK on enamel subsurface demineralization in vitro. Heliyon. 2024;10(1):e23176. doi:10.1016/j.heliyon.2023.e23176

7. Sakr AH, Nassif MS, El-Korashy DI. Amelogenin-inspired peptide, calcium phosphate solution, fluoride, and their synergistic effect on enamel biomimetic remineralization: an in vitro pH-cycling model. BMC Oral Health. 2024;24(1):279. doi:10.1186/s12903-024-04008-z

8. F. Torelli, Optogenetic control of odontoblastic differentiation in dental pulp stem cells (DPSC), Journal of Dental Sciences, 10.1016/j.jds.2026.01.021

9. Zheng W, Ding L, Wang Y, et al. The effects of 8DSS peptide on remineralization in a rat model of enamel caries evaluated by two nondestructive techniques. J Appl Biomater Funct Mater. 2019;17(1):1–8. doi:10.1177/2280800019827798

10. Shetty SS, Nekkanti S. Remineralization potential of a novel biomimetic material (self-assembling peptide P11-4) on early enamel caries: an in vitro study. J Contemp Dent Pract. 2023;24(3):181–187. doi:10.5005/jp-journals-10024-3490

11. Moura JS, Rodrigues LK, Del Bel Cury AA, Lima EM, Garcia RM. Influence of storage solution on enamel demineralization submitted to pH cycling. J Appl Oral Sci. 2004;12(3):205–208. doi:10.1590/S1678-77572004000300008

12. Hamdi K, Hamama HH, Motawea A, et al. Remineralization of early enamel lesions with a novel prepared tricalcium silicate paste. Sci Rep. 2022;12:9926. doi:10.1038/s41598-022-13608-0

13. Fu Y, Ekambaram M, Li KC, Zhang Y, Cooper PR, Mei ML. In vitro models used in cariology mineralisation research: a review of the literature. Dent J (Basel). 2024;12(10):323. doi:10.3390/dj12100323

14. Gouvêa DB, Santos NMD, Pessan JP, Jardim JJ, Rodrigues JA. Enamel subsurface caries-like lesions induced in human teeth by different solutions: a TMR analysis. Braz Dent J. 2020;31(2):157–163. doi:10.1590/0103-6440202003123

15. Hsu CC, Chung HY, Hagerman EM, Shi W, Yang JM, Wu B. Effects on hardness and elastic modulus for DSS-8 peptide treatment on remineralization of human dental tissues. Mater Res Soc Symp Proc. 2008;1132. doi:10.1557/PROC-1132-Z09-05

16. Kolumban A, Moldovan M, Ţig IA, Chifor I, Cuc S, Bud M, Badea ME. Evaluation of the demineralizing effects of various acidic solutions. Appl Sci. 2021;11:8270. doi:10.3390/app11178270

17. Duanis-Assaf T, Hu T, Lavie M, Zhang Z, Reches M. Understanding the adhesion mechanism of hydroxyapatite-binding peptide. Langmuir. 2022;38(3):968–978. doi:10.1021/acs.langmuir.1c02293

18. Argenta RM, Tabchoury CP, Cury JA. A modified pH-cycling model to evaluate fluoride effect on enamel demineralization. Pesqui Odontol Bras. 2003;17(3):241–246. doi:10.1590/S1517-74912003000300008

19. Borra RC, Lotufo MA, Gagioti SM, Barros FM, Andrade PM. A simple method to measure cell viability in proliferation and cytotoxicity assays. Braz Oral Res. 2009;23(3):255–262. doi:10.1590/S1806-83242009000300006

20. Pretty IA, Edgar WM, Higham SM. Detection of in vitro demineralization of primary teeth using quantitative light-induced fluorescence (QLF). Int J Paediatr Dent. 2002;12(3):158–167. doi:10.1046/j.1365-263X.2002.00357.x

21. Wu J, Donly ZR, Donly KJ, Hackmyer S. Demineralization depth using QLF and a novel image processing software. Int J Dent. 2010;2010:958264. doi:10.1155/2010/958264

22. Sezici YL, Çınarcık H, Yetkiner E, Attin R. Low-viscosity resin infiltration efficacy on postorthodontic white spot lesions: a quantitative light-induced fluorescence evaluation. Turk J Orthod. 2020;33(2):92–97. doi:10.5152/TurkJOrthod.2020.19088

23. Gmür R, Giertsen E, van der Veen MH, de Josselin de Jong E, ten Cate JM, Guggenheim B. In vitro quantitative light-induced fluorescence to measure changes in enamel mineralization. Clin Oral Investig. 2006;10(3):187–195. doi:10.1007/s00784-006-0058-z

24. Beltrami R, Colombo M, Rizzo K, et al. Cytotoxicity of different composite resins on human gingival fibroblast cell lines. Biomimetics (Basel). 2021;6(2):26. doi:10.3390/biomimetics6020026

25. Gomez J. Detection and diagnosis of the early caries lesion. BMC Oral Health. 2015;15(Suppl 1):S3. doi:10.1186/1472-6831-15-S1-S3

26. Bang J, Park H, Yoo J, et al. Selection and identification of a novel bone-targeting peptide for biomedical imaging of bone. Sci Rep. 2020;10:10576. doi:10.1038/s41598-020-67522-4

27. Aydın B, Pamir T, Baltaci A, Orman MN, Turk T. Effect of storage solutions on microhardness of crown enamel and dentin. Eur J Dent. 2015;9(2):262–266. doi:10.4103/1305-7456.156848

28. Hardan L, Chedid JCA, Bourgi R, et al. Peptides in dentistry: a scoping review. Bioengineering (Basel). 2023;10(2):214. doi:10.3390/bioengineering10020214

29. Zawawi R, Almosa N. Cariogenic enamel demineralization prevention, detection, and management: a literature review. Eur J Dent. 2025. doi:10.1055/s-0045-1809179

30. Kunert M, Rozpedek-Kaminska W, Galita G, et al. The cytotoxicity and genotoxicity of bioactive dental materials. Cells. 2022;11(20):3238. doi:10.3390/cells11203238

31. Gegamyan AO, Sarap LR, Zeibert AY. Evaluation of enamel remineralization rate by quantitative light-induced fluorescence. Clin Dent (Russ). 2021;24(4):13–17. doi:10.37988/1811-153X_2021_4_13

32. Long JR, Dindot JL, Zebroski H, et al. A peptide that inhibits hydroxyapatite growth is in an extended conformation on the crystal surface. Proc Natl Acad Sci U S A. 1998;95(21):12083–12087. doi:10.1073/pnas.95.21.12083

33. Torelli F. Visible-light-triggered BMP-2 release from enzymatically crosslinked marine collagen-alginate hydrogel blends enhances osteogenesis in dental pulp stem cells. Front Physiol. 2026 Feb 17;17:1743209. doi:10.3389/fphys.2026.1743209.

34. Townsend L, Williams RL, Anuforom O, et al. Antimicrobial peptide coatings for hydroxyapatite: electrostatic and covalent attachment of antimicrobial peptides to surfaces. J R Soc Interface. 2017;14(130):20160657. doi:10.1098/rsif.2016.0657

35. Weiger MC, Park JJ, Roy MD, Stafford CM, Karim A, Becker ML. Quantification of the binding affinity of a specific hydroxyapatite-binding peptide. Biomaterials. 2010;31(11):2955–2963. doi:10.1016/j.biomaterials.2010.01.012

36. Ayoub HM, Gregory RL, Tang Q, Lippert F. Influence of salivary conditioning and sucrose concentration on biofilm-mediated enamel demineralization. J Appl Oral Sci. 2020;28:e20190501. doi:10.1590/1678-7757-2019-0501

37. Enax J, Fandrich P, Schulze zur Wiesche E, Epple M. The remineralization of enamel from saliva: a chemical perspective. Dent J (Basel). 2024;12(11):339. doi:10.3390/dj12110339

38. Enax J, Ganss B, Amaechi BT, Meyer F. The composition of the dental pellicle: an updated literature review. Front Oral Health. 2023;4:1260442. doi:10.3389/froh.2023.1260442

39. Ferrari-Peron P, Steuer LM, Schmidtmann I, et al. In vivo comparison of initial caries lesions using the enamel decalcification index and quantitative light-induced fluorescence during orthodontic therapy. Clin Oral Investig. 2025;29(3):174. doi:10.1007/s00784-025-06234-3

40. Hegde MN, Sajnani AR. Salivary proteins: a barrier on enamel demineralization—an in vitro study. Int J Clin Pediatr Dent. 2017;10(1):10–13. doi:10.5005/jp-journals-10005-1398

41. Marin LM, Xiao Y, Cury JA, Siqueira WL. Engineered salivary peptides reduce enamel demineralization provoked by cariogenic Streptococcus mutans biofilm. Microorganisms. 2022;10(4):742. doi:10.3390/microorganisms10040742

42. Roberts JM, Bradshaw DJ, Lynch RJM, Higham SM, Valappil SP. Quantifying demineralisation of enamel using a hyperspectral camera measuring fluorescence loss. Photodiagnosis Photodyn Ther. 2021;36:102603. doi:10.1016/j.pdpdt.2021.102603

43. Ando M, van der Veen MH, Schemehorn BR, Stookey GK. Comparative study to quantify demineralized enamel in deciduous and permanent teeth using laser-and light-induced fluorescence techniques. Caries Res. 2001;35(6):464–470. doi:10.1159/000047491

44. Xiao Q, Tu R, He T, et al. Evaluation of fluorescence imaging with reflectance enhancement (FIRE) for quantifying enamel demineralization in vitro. Caries Res. 2015;49(5):531–539. doi:10.1159/000365298

45. Lavis LD, Raines RT. Bright ideas for chemical biology. ACS Chem Biol. 2008;3(3):142–155. doi:10.1021/cb700248m

